# Functional diversity increases the efficacy of phage combinations

**DOI:** 10.1101/2021.07.09.451746

**Authors:** Rosanna C. T. Wright, Ville-Petri Friman, Margaret C. M. Smith, Michael A. Brockhurst

**Affiliations:** Faculty of Biology, Medicine and Health, University of Manchester, Manchester, UK; Department of Biology, University of York, York, UK

## Abstract

Phage therapy is a promising alternative to traditional antibiotics for treating bacterial infections. Such phage-based therapeutics typically contain multiple phages, but how the efficacy of phage combinations scales with phage richness, identity and functional traits is unclear. Here, we experimentally tested the efficacy of 827 unique phage combinations ranging in phage richness from 1 to 12 phages. The efficacy of phage combinations increased with phage richness. However, complementarity between functionally diverse phages allowed efficacy to be maximised at lower levels of phage richness in functionally diverse combinations. These findings suggest that phage functional diversity is the key property of effective phage combinations, enabling the design of simple but effective phage therapies that overcome the practical and regulatory hurdles that limit development of more diverse phage therapy cocktails.

## Introduction

Bacterial killing by lytic phages regulates bacterial turnover in microbial communities, influencing bacterial community dynamics in both environmental and clinical settings [1, 2]. Phage diversity is predicted to exceed that of their hosts by up to 10 times [3], and may vary between communities just metres apart [4]. A survey of phages able to infect *Pseudomonas aeruginosa*, a commonly multi-drug resistant opportunistic pathogen [5], across 4 continents identified 7 distinct phage groups with lytic activity against 87% of clinical bacterial strains tested [6], and novel phage taxa are continually being discovered [7, 8]. This diversity of phages offers a promising alternative to antibiotics for treating bacterial infection where rates of antibiotic resistance are rapidly rising [9–11]. Multiple phages are often combined for therapeutic use to improve the range of hosts which can be targeted. However, more diverse combinations pose greater regulatory hurdles as individual phages and interaction effects must be assessed [12–14]. While developing efficient, low diversity phage combinations would be highly practical, it is unclear which rationale one should use to design such combinations.

The relationship between biodiversity and ecosystem functioning (i.e., the collective activity of a community) is usually positive [15]. Increasing species richness has been shown to improve the function of microbial communities by two distinct mechanisms [16, 17]. Firstly, if species perform different ecological roles, then greater species richness can deliver higher community-level performance due to functional complementarity, through filling more of the available niche space [16, 18]. Secondly, more diverse communities are more likely to contain highly performing taxa simply by chance, leading to a positive relationship between diversity and function due to species identity effects [16, 17]. On the other hand, functional redundancy among species in a community can lead to diminishing returns of further increasing species richness, resulting in a saturating relationship between richness and function [19–21]. These counteracting effects suggest that high phage efficacy could be attained by low richness phage combinations, provided that these contain functionally different and non-redundant phages. Such combinations would have practical benefits in terms of reducing the manufacturing and regulatory challenges posed by higher order phage combination therapies [13, 14, 22].

For phages, lytic infection of a bacterial host depends on adsorption to the host outer membrane and evasion of host phage defence systems once within the cell [23]. Binding to specific bacterial cell-surface receptors for adsorption is therefore a key functional trait for phages. Phage combinations targeting higher numbers of receptors could be considered functionally diverse and more efficacious by reducing competition among phages for shared adsorption sites. Such phage combinations could also limit resistance evolution via cell surface modification since this would likely require multiple mutations [24–26], that often impose additive fitness costs [27].

By applying the principles of biodiversity-ecosystem functioning to the design of phage therapies, we sought to determine the relative importance of phage species richness, functional diversity and identity effects on the efficacy of phage combinations. We used twelve *Pseudomonas aeruginosa* phages including phages targeting either lipopolysaccharide (LPS) or Type IV pilus (T4P) for adsorption to result in 827 unique phage combinations with differing levels of species richness and degrees of functional diversity. These combinations included all possible single, pairwise and 3-member communities, 264 different 4- and 6-member communities and the full 12-member community. We show that phage richness had a saturating relationship with efficacy (defined as the suppression of bacterial growth), and that highly efficacious but low richness phage combinations could be designed provided that they had high functional diversity, *i.e.*, the constituent phages targeted multiple distinct adsorption receptors. Together, these results suggest that ecological complementarity plays a key role in determining the efficacy of phage combinations.

## Materials and methods

### Phage combination design and community assembly

A panel of 12 lytic *P. aeruginosa* phages were used to build phage combinations of varying phage community species richness. The adsorption receptors of all phages has previously been characterised [28]; 4 phages adsorb via T4P, and 8 via LPS. In total, we assembled 827 different phage combinations, ranging from single phage (12), all possible 2- and 3-member communities (66 and 220 respectively), a random partition of 4- and 6-member communities (264 of each), to the full 12-member community. A random partition design was used to select 4 and 6-phage communities which equally represent all phage strains across both richness levels (as described previously [29]).

Phage stocks were amplified to equal densities (~8.9×10^10^±1.0×10^11^ pfu/ml) using the susceptible bacterial host *P. aeruginosa* PAO1, isolated by filtration (0.22μm) and stored at 4°C. To limit human error, master plates of phage communities were assembled in deep 96 well plates using a liquid handling robot (epMotion^®^ 5070, Eppendorf, Germany) in triplicate. Equal volumes (and densities) of each phage were added to give a final volume of 120μl per phage community.

### Measuring the efficacy of phage combinations

We determined efficacy of phage combinations as their ability to supress growth of the susceptible host, *P. aeruginosa* PAO1. Bacterial replicates were inoculated from three single colonies into 6ml KB and grown overnight at 37°C with shaking at 180rpm, before diluting 100-fold into assay plates containing 120μl of KB. Phage communities were transferred from the master plates (15μl per well) to give a multiplicity of infection of approximately 100 phage per bacterial cell (actual MOI ~80.4±20.2 with initial bacterial density of ~9.6×10^7^±1.1×10^7^). Optical density (absorbance at 600nm; *Abs*_600_) was measured immediately, then after static 24h incubation at 37°C. Phage combination efficiency was measured as reduction in bacterial growth in the presence relative to the absence of phage, as ‘Efficacy’ (Eq 1; [30]).

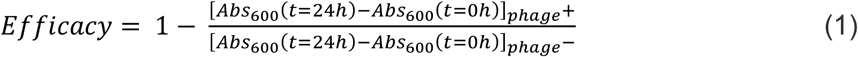

For each phage combination, the phage with the highest independent Efficacy value was considered as the best constituent phage (*i.e.*, measured when Diversity = 1; matched by replicate to the phage combination). These values were used to calculate transgressive overyielding, D_max_ (Eq. 2; [31]), which is a measure that describes the efficacy of a community relative to its most efficient member, such that a value of 0 indicates equal efficacy and values above 0 indicates that the community was more effective.

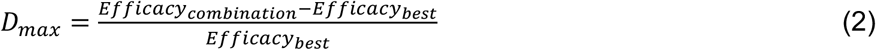

### Statistical analysis

The relationship between phage strain richness and efficacy was analysed using a previously described linear model method designed to separate significant factors affecting biodiversity-ecosystem functioning [29]. Phage richness and functional diversity (*i.e.*, number of different receptor targets) were included as interacting main effects, alongside other main effects of receptor targets (*i.e.*, presence of LPS and/or T4P targeting phages), phage identity (including pairwise and higher order interactions between phages as separate main effects) and non-linear effects of phage richness. Due to the saturating relationship between phage richness and efficacy, we also fitted a non-linear asymptotic exponential model to the data. Model parameters were determined using the nonlinear least squares function in R [32], and compared to equivalent linear models using Akaike Information Criterion (AIC).

The effects of richness, functional diversity and phage identity on transgressive overyielding were also analysed using a linear model. Significant interaction terms between functional diversity and richness, functional diversity and phage identity, and pairwise interactions between phages were included in the model. Here, receptor target was excluded from the model as it was not significant. Non-linear richness was included as an explanatory variable, and best fit of regression models (linear versus decaying exponential model) were assessed by AIC to fit regression curves to the plotted data. All analyses were performed in R (version 3.5.2; [32]).

## Results

### Diminishing returns of increasing phage richness on phage combination efficacy

The efficacy of phage combinations, measured as their ability to reduce bacterial growth, increased with phage community richness (*i.e.*, number of phage strains). However, phage richness explained only 30% of variation in efficacy (Figure 1; linear model_efficacy_: F_1,5740_=2497, p<0.0001, R^2^=0.303). Non-linear models explained a greater proportion of the variation: the relationship between richness and efficacy was best explained by an asymptotic exponential model where the asymptote was reached when bacterial growth was completely suppressed (Figure 1; AIC _linear model_ = 205; AIC _asymptotic model_ = −1190). This suggests that there were low richness phage combinations that were as effective as the highest richness phage combination, whose efficiency could not be improved with additional phages, likely due to functional redundancy among the constituent phage strains.

**Figure 1.**
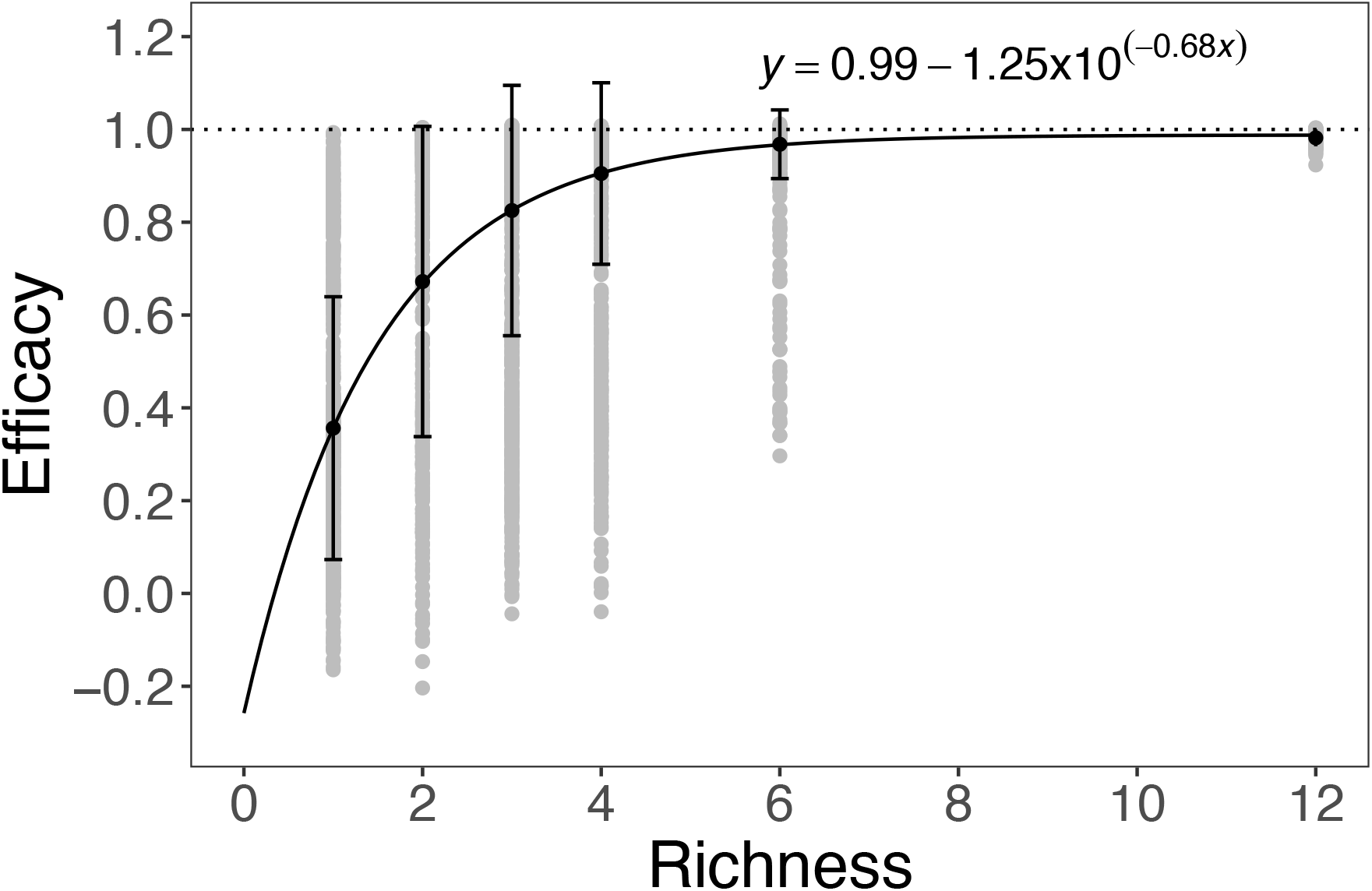
Saturating relationship between the efficacy and richness of phage combinations. Efficiency of phage combinations measured as mean efficacy (± s.d.) of bacterial growth suppression in the presence of phage relative to phage-free growth, raw data in grey. The dashed line at 1 indicates complete suppression of bacterial growth by the phage community. An asymptotic exponential with the equation shown was fit to the data using a non-linear least squares model.

### Increased phage functional diversity improves phage combination efficacy

Including functional diversity (FD), in terms of the receptor targeted by the phage, in the model explained substantially more of the observed variation in efficacy: phage functional diversity and strain richness together accounted for ~70% of the variation in efficacy (Figure 2; linear model_efficacy_, main effects of phage diversity and functional diversity and their interaction: F_3,5738_=4476, p<0.0001, R^2^=0.701). Phages targeting T4P contributed more to efficacy than LPS-binding phages, such that combinations containing only T4P-binding phages outperformed combinations containing only LPS-binding phages (Figure 2A+B; linear model_efficacy_, coefficient of T4P-binding phages: t=11.16, p<0.0001; coefficient of LPS-binding phages: t=(−)24.57, p<0.001). Functionally diverse combinations containing phages targeting both LPS and T4P receptors showed a saturating relationship between efficacy and phage richness and completely supressed bacterial growth at lower levels of phage diversity than combinations targeting only a single receptor (Figure 2C). Phage identity accounted for only ~3% of the remaining variation in efficacy (linear model_efficacy_: F_12,5730_=15.45, p<0.0001). Consistent with stronger complementarity effects at higher functional diversity, we observed greater transgressive overyielding for high compared to low functional diversity combinations (Figure 3; linear model_Dmax_, F_1,5740_=498.7, p<0.0001). Moreover, transgressive overyielding was stronger at lower phage richness levels for high but not low functional diversity combinations (Figure 3, linear model_Dmax_, richness F_1,5740_=31.1, p<0.0001; interaction: functional diversity * richness F_3,5738_=136, p<0.0001). Together these data suggest that although the presence of certain phage strains could influence efficacy, functional diversity was the strongest predictor, leading to high efficacy even in low richness phage combinations.

**Figure 2.**
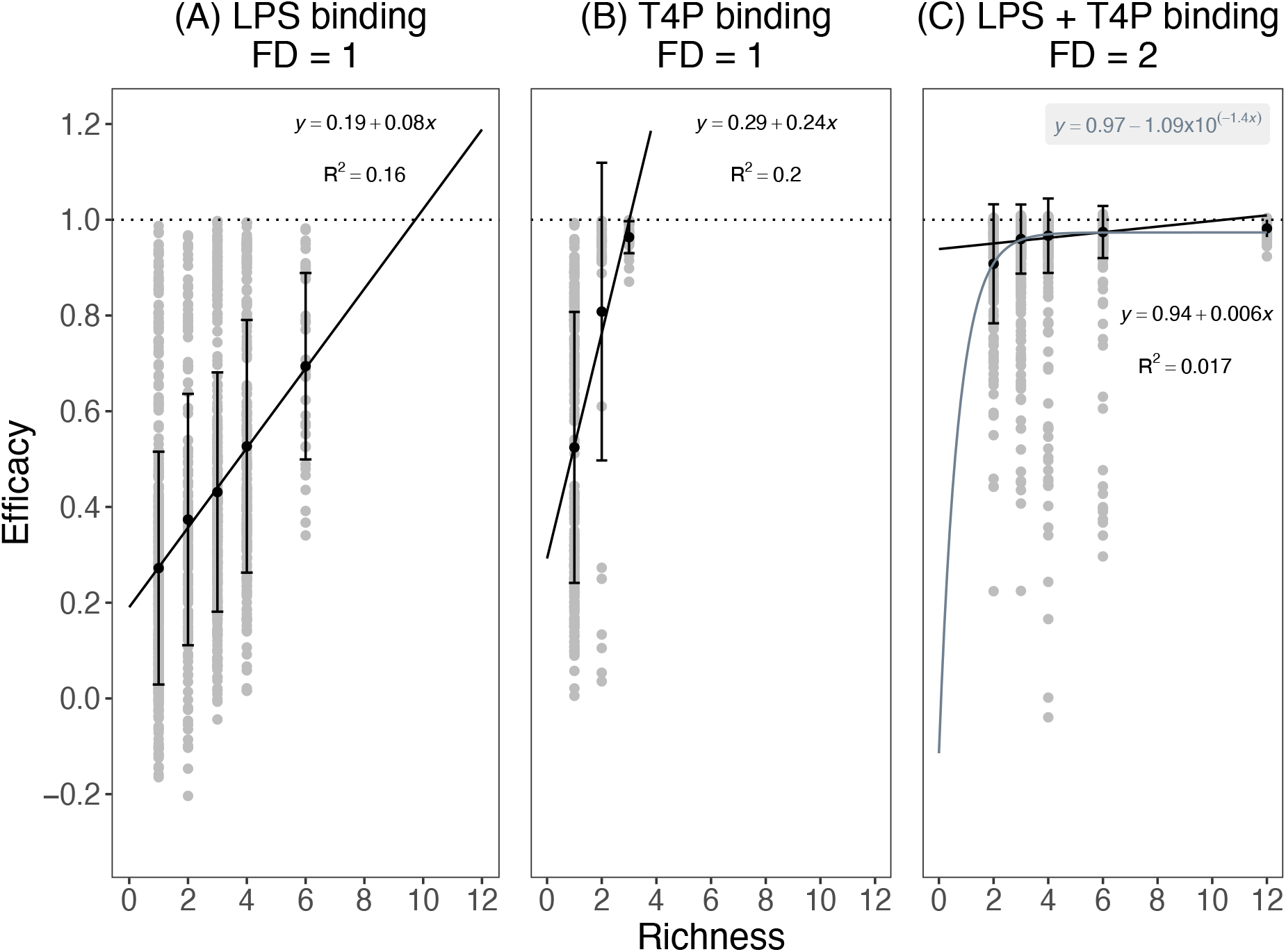
Functional diversity increases efficacy of phage combinations. Efficacy of phage combinations with low functional diversity (FD = 1), where all phage target either LPS (A) or Type IV pilus (B), and phage combinations with high functional diversity (FD = 2) which include phages targeting both the LPS and Type IV Pilus (C). Efficacy is measured as suppression of bacterial growth by phages relative to phage-free populations; mean values (± s.d.) are shown in black, with raw data in grey to show distribution of data points. The dashed line indicates the theoretical maximum reduction (*i.e.*, no bacterial growth detected). Linear regression equations relate to relationship between phage richness (x) and efficacy (y); note that for (C), an asymptotic exponential model (shown in grey) better explains this relationship (AIC _linear model_ = 205; AIC _asymptotic model_ = −1190).

**Figure 3.**
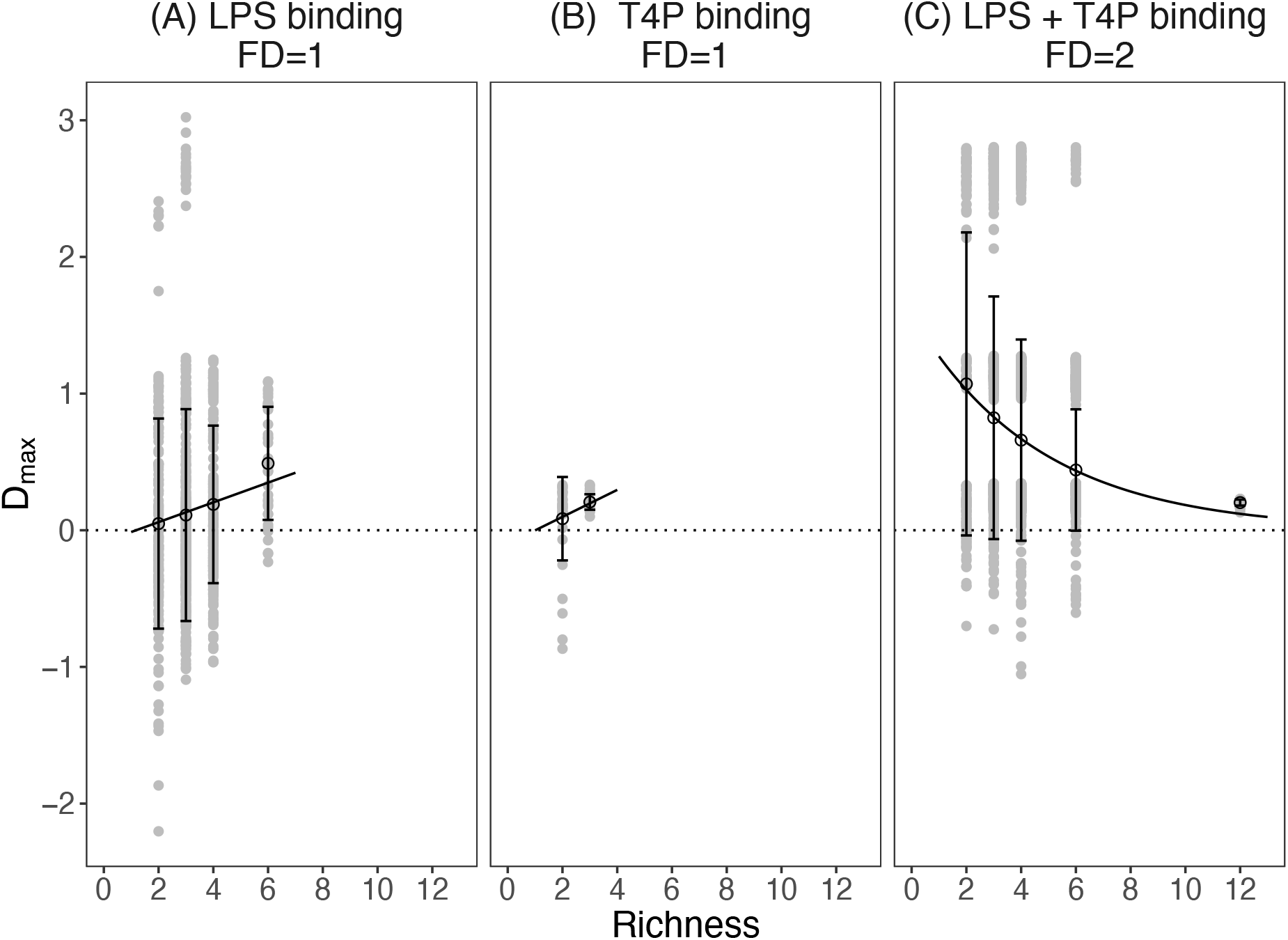
Degree of transgressive overyielding is determined by phage functional diversity. Transgressive overyielding, D_max_, describes the efficacy of phage combinations relative to the best constituent phage as a monoculture. Phage combinations either target one receptor (FD = 1; A, LPS binding; B, T4P binding) or include phages targeting LPS and targeting T4P (FD = 2; C, LPS + T4P binding). Mean values (± s.d.) are shown in black, with raw data in grey to show distribution of data points; regression lines were fit either as a linear model (A-B) or an exponential degradation model (C).

## Discussion

To enable rational design of phage therapy combinations, it is important to understand the key factors which determine a phage combination’s efficacy in supressing bacterial growth. Applying concepts from the analysis of ecological biodiversity-ecosystem function relationships, we compared the relative contributions of phage richness, phage identity and functional diversity in determining the efficacy of phage combinations. We observed a saturating relationship between phage richness and efficacy, consistent with diminishing returns of increasing richness due to functional redundancy among phages at higher richness levels. Correspondingly, phage combinations with higher functional diversity, in terms of the number of cell surface receptors targeted for phage adsorption, were more effective at suppressing bacterial growth and were able to do so at lower levels of phage richness (e.g., 2-3 phages) than low functional diversity combinations. Functionally diverse phage combinations targeting different adsorption receptors displayed higher transgressive overyielding and stronger complementarity at low levels of phage richness, achieving up to 3-fold higher efficacy than their best constituent phage even for combinations of just two phages. Together, our data suggest that functional diversity is the most important determinant of the efficacy of phage combinations.

By maximising functional diversity, phage combinations can be optimised for bacterial killing at low strain diversity, thus reducing the regulatory hurdles of preparing more complex therapeutic combinations [13, 14, 22]. Functional complementarity between phages targeting different adsorption receptors is likely to have two key benefits: Firstly, decreased competition for binding sites to adsorb to the bacterial cell may lead to increased lysis. Secondly, functionally diverse phage combinations are more likely to suppress resistance evolution. The majority of resistance mutations arising against our phage panel target the genes encoding the bacterial cell surface receptors (LPS and T4P; [28, 33]), and as such, promote cross-resistance to alternative phages which adsorb to the same receptor. In contrast, resistance to a functionally diverse phage combination is likely require multiple independent resistance mutations (e.g. modification of each adsorption target; [33]) which will co-occur in the same cell with far lower probability.

In addition to increasing efficacy against a single bacterial genotype, higher functional diversity may also prove beneficial in more complex scenarios. For example, functionally diverse phage combinations are likely to be able to target a broader diversity of bacterial genotypes. This could be particularly relevant in treatment of chronic infections, where the bacterial populations typically undergo extensive evolutionary diversification (e.g., in response to host-pathogen interactions) [34, 35]. This can lead to altered expression of common phage receptor targets, including modification and even loss of LPS components and T4P [36–38], which can reduce susceptibility to phage infection [39, 40]. Essentiality of different cell-surface receptors across environments may explain differences in observed efficacy between phages targeting different adsorption receptors. Phage combinations targeting a broader range of cell surface receptors will be more likely to be able to infect and clear such host-adapted bacterial populations.

In this study, functional phage diversity was limited to two cell surface receptor targets, but further increases in the diversity of receptors targeted by phage combinations is likely to lead to further increases in their efficacy. Examples of other *P. aeruginosa* cell surface receptors used for phage adsorption include, outer membrane porins [41] and other membrane anchored proteins such as TonB-dependent receptors, which can be involved in iron-siderophore uptake [24, 42]. An additional limitation of this study is that we did not test a strain encoding a CRISPR-Cas immunity system [43]. Inducible resistance mechanisms may be preferentially selected *in vivo* because of their lower fitness costs compared to surface receptor modification mutations [44]. Unlike surface modification resistance mutations, CRISPR-mediated resistance is likely to promote different cross-resistance interactions between phages mediated by their genetic similarity rather than their receptor target for adsorption. This suggests that whilst functional diversity of phage strains is necessary to limit the evolution of cross-resistance via surface modification, maximising genetic diversity could be important to limit cross-resistance via sequence-based resistance mechanisms such as CRISPR-Cas, restriction modification or other recently discovered phage defence systems [45].

To conclude, our findings suggest that maximising functional diversity is a simple and effective rule for designing high efficacy, low richness phage combinations overcoming the regulatory hurdles associated with preparation of complex phage cocktails.

## Abbreviations

T4P: Type IV Pilus
LPS: Lipopolysaccharide

## Acknowledgements

This work was supported by an ACCE DTP PhD Studentship to RCTW funded by the Natural Environment Research Council (NE/L002450/1, Studentship 1517986). We thank Matti Jalasvuori for providing us with phage strains.

